# Loss of cytoplasmic incompatibility and minimal fecundity effects explain relatively low *Wolbachia* frequencies in *Drosophila mauritiana*

**DOI:** 10.1101/461574

**Authors:** Megan K. Meany, William R. Conner, Sophia V. Richter, Jessica A. Bailey, Michael Turelli, Brandon S. Cooper

**Author notes:** Correspondence: Division of Biological Sciences, University of Montana, 32 Campus Drive, ISB, Missoula, MT 59812, USA.

## Abstract

Maternally transmitted *Wolbachia* bacteria infect about half of all insect species. Many *Wolbachia* cause cytoplasmic incompatibility (CI), reduced egg hatch when uninfected females mate with infected males. Although CI produces a frequency-dependent fitness advantage that leads to high equilibrium *Wolbachia* frequencies, it does not aid *Wolbachia* spread from low frequencies. Indeed, the fitness advantages that produce initial *Wolbachia* spread and maintain non-CI *Wolbachia* remain elusive. *w*Mau *Wolbachia* infecting *Drosophila mauritiana* do not cause CI, despite being very similar to CI-causing *w*No from *D. simulans* (0.068% sequence divergence over 682,494 bp), suggesting recent CI loss. Using draft *w*Mau genomes, we identify a deletion in a CI-associated gene, consistent with theory predicting that selection within host lineages does not act to increase or maintain CI. In the laboratory, *w*Mau shows near-perfect maternal transmission; but we find no significant effect on host fecundity, in contrast to published data. Intermediate *w*Mau frequencies on the island Mauritius are consistent with a balance between unidentified small, positive fitness effects and imperfect maternal transmission. Our phylogenomic analyses suggest that group-B *Wolbachia*, including *w*Mau and *w*Pip, diverged from group-A *Wolbachia*, such as *w*Mel and *w*Ri, 6–46 million years ago, more recently than previously estimated.

## INTRODUCTION

Maternally transmitted *Wolbachia* infect about half of all species throughout all major insect orders (Werren and Windsor 2000; Zug and Hammerstein 2012; Weinert et al. 2015), as well as other arthropods (Jeyaprakash and Hoy 2000; Hilgenboecker et al. 2008) and nematodes (Taylor et al. 2013). Host species may acquire *Wolbachia* from common ancestors, from sister species via hybridization and introgression, or horizontally (O’Neill et al. 1992; Rousset and Solignac 1995; Huigens et al. 2004; Baldo et al. 2008; Raychoudhury et al. 2009; Gerth and Bleidorn 2016; Schuler et al. 2016; Turelli et al. 2018). *Wolbachia* often manipulate host reproduction, inducing cytoplasmic incompatibility (CI) and male killing in *Drosophila* (Laven 1951; Yen and Barr 1971; Hoffmann et al. 1986; Hoffmann and Turelli 1997; Hurst and Jiggins 2000). CI reduces egg hatch when *Wolbachia*-uninfected females mate with infected males. Three parameters usefully approximate the frequency dynamics and equilibria of CI-causing *Wolbachia* that do not distort sex ratios: the relative hatch rate of uninfected eggs fertilized by infected males (*H*), the fitness of infected females relative to uninfected females (*F*), and the proportion of uninfected ova produced by infected females (*μ*) (Caspari and Watson 1959; Hoffmann et al. 1990). To spread deterministically from low frequencies, *Wolbachia* must produce *F*(1 – *μ*) > 1, irrespective of CI. Once they become sufficiently common, CI-causing infections, such as *w*Ri-like *Wolbachia* in *Drosophila simulans* and several other *Drosophila* (Turelli et al. 2018), spread to high equilibrium frequencies, dominated by a balance between CI and imperfect maternal transmission (Turelli and Hoffmann 1995; Kreisner et al. 2016). In contrast, non-CI-causing *Wolbachia*, such as *w*Au in *D. simulans* (Hoffmann et al. 1996), typically persist at lower frequencies, presumably maintained by a balance between positive *Wolbachia* effects on host fitness and imperfect maternal transmission (Hoffmann and Turelli 1997; Kreisner et al. 2013). When *H* < *F*(1 – *μ*) < 1, “bistable” dynamics result, producing stable equilibria at 0 and at a higher frequency denoted *p*_*s*_, where 0.50 < *p*_*s*_ ≤ 1 (Turelli and Hoffmann 1995). Bistability explains the pattern and (slow) rate of spread of *w*Mel transinfected into *Aedes aegypti* to suppress the spread of dengue, Zika and other human diseases (Hoffmann et al. 2011; Barton and Turelli 2011; Turelli and Barton 2017; Schmidt et al. 2017).

In contrast to the bistability observed with *w*Mel transinfections, natural *Wolbachia* infections seem to spread via “Fisherian” dynamics with *F*(1 – *μ*) > 1 (Fisher 1937; Kriesner et al. 2013; Hamm et al. 2014). Several *Wolbachia* effects could generate *F*(1 – *μ*) > 1, but we do not yet know which ones actually do. For example, *w*Ri has evolved to increase *D. simulans* fecundity in only a few decades (Weeks et al. 2007), *w*Mel seems to enhance *D. melanogaster* fitness in high and low iron environments (Brownlie et al. 2009), and several *Wolbachia* including *w*Mel protect their *Drosophila* hosts from RNA viruses (Hedges et al. 2008; Teixeira et al. 2008; Martinez et al. 2014). However, it remains unknown which if any these potential fitness benefits underlie *Wolbachia* spread in nature. For instance, *w*Mel seems to have little effect on viral abundance in wild-caught *D. melanogaster* (Webster et al. 2015; Shi et al. 2018).

*D. mauritiana, D. simulans* and *D. sechellia* comprise the *D. simulans* clade within the nine-species *D. melanogaster* subgroup of *Drosophila*. The *D. simulans* clade diverged from *D. melanogaster* approximately three million years ago (mya), with the island endemics *D. sechellia* (Seychelles archipelago) and *D. mauritiana* (Mauritius) thought to originate in only the last few hundred thousand years (Lachaise et al. 1986; Ballard 2000a; Dean and Ballard 2004; McDermott and Kliman 2008; Garrigan et al. 2012; Brand et al. 2013; Garrigan et al. 2014). *D. simulans* is widely distributed around the globe, but has never been collected on Mauritius (David et al. 1989; Legrand et al. 2011). However, evidence of mitochondrial and nuclear introgression supports interisland migration and hybridization between these species (Ballard 2000a; Nunes et al. 2010; Garrigan et al. 2012), which could allow introgressive *Wolbachia* transfer (Rousset and Solignac 1995).

*D. mauritiana* is infected with *Wolbachia* denoted *w*Mau, likely acquired via introgression from other *D. simulans*-clade hosts (Rousset and Solignac 1995). *Wolbachia* variant *w*Mau may also infect *D. simulans* (denoted *w*Ma in *D. simulans*) in Madagascar and elsewhere in Africa and the South Pacific (Ballard 2000a; Ballard 2004). *w*Mau does not cause CI in *D. mauritiana* or when transinfected into *D. simulans* (Giordano et al. 1995). Yet it is very closely related to *w*No strains that do cause CI in *D. simulans* (Merçot et al. 1995; Rousset and Solignac 1995; James and Ballard 2000, 2002). (Also, *D. simulans* seems to be a “permissive” host for CI, as evidenced by the fact that *w*Mel, which causes little CI in its native host, *D. melanogaster*, causes intense CI in *D. simulans* [Poinsot et al. 1998].) Fast et al. (2011) reported that a *w*Mau variant increased *D. mauritiana* fecundity four-fold. This fecundity effect occurred in concert with *w*Mau-induced alternations of programmed cell death in the germarium and of germline stem cell mitosis, possibly providing insight into the mechanisms underlying increased egg production (Fast et al. 2011). However, the generality of this finding across *w*Mau variants and host genetic backgrounds remains unknown.

Here, we assess the genetic and phenotypic basis of *w*Mau frequencies in *D. mauritiana* on Mauritius by combining analysis of *w*Mau draft genomes with analysis of *w*Mau transmission in the laboratory and *w*Mau effects on host fecundity and egg hatch. We identify a single mutation that disrupts a locus associated with CI. The loss of CI in *w*Mau is consistent with theory demonstrating that selection within host species does not act to increase or maintain the level of CI (Prout 1994; Turelli 1994; Haygood and Turelli 2009), but instead acts to increase *F*(1 – *μ*), the product of *Wolbachia* effects on host fitness and maternal transmission efficiency (Turelli 1994). The loss of CI helps explain the intermediate *w*Mau frequencies on Mauritius, reported by us and Giordano et al. (1995). We find no *w*Mau effects on host fecundity, and theoretical analyses show that even a two-fold fecundity increase cannot be reconciled with the observed intermediate population frequencies, unless maternal *w*Mau transmission is exceptionally unreliable in the field. Finally, we present theoretical analyses illustrating that the persistence of two distinct classes of mtDNA haplotypes among *Wolbachia*-uninfected *D. mauritiana* is unexpected under a simple null model. Together, our results contribute to understanding the genomic and phenotypic basis of global *Wolbachia* persistence, which is relevant to improving *Wolbachia*-based biocontrol of human diseases (Ritchie 2018).

## MATERIALS AND METHODS

### *Drosophila* Husbandry and Stocks

The *D. mauritiana* isofemale lines used in this study (*N* = 32) were sampled from Mauritius in 2006 by Margarita Womack and kindly provided to us by Prof. Daniel Matute from the University of North Carolina, Chapel Hill. We also obtained four *D. simulans* stocks (lines 196, 297, 298, and 299) from the National *Drosophila* Species Stock Center that were sampled from Madagascar. Stocks were maintained on modified version of the standard Bloomington-cornmeal medium (Bloomington Stock Center, Bloomington, IN) and were kept at 25°C, 12 light:12 dark photoperiod prior to the start of our experiments.

### Determining *Wolbachia* infection status and comparing infection frequencies

One to two generations prior to our experiments DNA was extracted from each isofemale line using a standard ‘squish’ buffer protocol (Gloor et al. 1993), and infection status was determined using a polymerase chain reaction (PCR) assay (Simpliamp ThermoCycler, Applied Biosystems, Singapore). We amplified the *Wolbachia*-specific *wsp* gene (Forward: 5’-TGGTCCAATAAGTGATGAAGAAAC-3’; Reverse: 5’-AAAAATTAAACGCTACTCCA-3’; Braig et al. 1998) and a nuclear control region of the *2L* chromosome (Forward: 5’-TGCAGCTATGGTCGTTGACA-3’; Reverse: 5’-ACGAGACAATAATATGTGGTGCTG-3’; designed here). PCR products were visualized using 1% agarose gels that included a molecular-weight ladder. Assuming a binomial distribution, we estimated exact 95% binomial confidence intervals for the infection frequencies on Mauritius. Using Fisher’s Exact Test, we tested for temporal differences in *w*Mau frequencies by comparing our frequency estimate to a previous estimate (Giordano et al. 1995). All analyses were performed using R version 3.5.1 (R Team 2015).

We used quantitative PCR (qPCR) (MX3000P, Agilent Technologies, Germany) to confirm that tetracycline-treated flies were cleared of *w*Mau. DNA was extracted from *D. mauritiana* flies after four generations of tetracycline treatment (1-2 generations prior to completing our experiments), as described below. Our qPCR used a PowerUp™ SYBR™ Green Master Mix (Applied Biosystems™, California, USA) and amplified *Wolbachia*-specific *wsp* (Forward: 5’-CATTGGTGTTGGTGTTGGTG-3’; Reverse: 5’-ACCGAAATAACGAGCTCCAG-3’) and *Rpl32* as a nuclear control (Forward: 5’-CCGCTTCAAGGGACAGTATC-3’; Reverse: 5’-CAATCTCCTTGCGCTTCTTG-3’; Newton and Sheehan 2014).

### *Wolbachia* DNA extraction, library preparation, and sequencing

We sequenced *w*Mau-infected *R9, R29*, and *R60 D. mauritiana* genotypes. Tissue samples for genomic DNA were extracted using a modified CTAB Genomic DNA Extraction protocol. DNA quantity was tested on an Implen Nanodrop (Implen, München, Germany) and total DNA was quantified by Qubit Fluorometric Quantitation (Invitrogen, Carlsbad, California, USA). DNA was cleaned using Agencourt AMPure XP beads (Beckman Coulter, Inc., Brea, CA, U.S.A), following manufacturers’ instructions, and eluted in 50 μl 1× TE Buffer for shearing. DNA was sheared using a Covaris E220 Focused Ultrasonicator (Covaris Inc., Woburn, MA) to a target size of 400 bp. We prepared libraries using NEBNext® Ultra™ II DNA Library Prep with Sample Purification Beads (New England BioLabs, Ipswich, Massachusetts). Final fragment sizes and concentrations were confirmed using a TapeStation 2200 system (Agilent, Santa Clara, California). We indexed samples using NEBNext® Multiplex Oligos for Illumina® (Index Primers Set 3 & Index Primers Set 4), and 10 μl of each sample was shipped to Novogene (Sacramento, CA) for sequencing using Illumina HiSeq 4000 (San Diego, CA), generating paired-end, 150 bp reads.

### *Wolbachia* assembly

We obtained published reads (*N* = 6) from Garrigan et al. (2014), and assembled these genomes along with the *R9, R29*, and *R60* genomes that we sequenced. Reads were trimmed using Sickle v. 1.33 (Joshi and Fass 2011) and assembled using ABySS v. 2.0.2 (Jackman et al. 2017). *K* values of 41, 51, 61, and 71 were used, and scaffolds with the best nucleotide BLAST matches to known *Wolbachia* sequences with E-values less than 10^−10^ were extracted as the draft *Wolbachia* assemblies. We deemed samples infected if the largest *Wolbachia* assembly was at least 1 million bases and uninfected if the largest assembly was fewer than 100,000 bases. No samples produced *Wolbachia* assemblies between 100,000 and 1 million bases. Of the six sets of published reads we analyzed (Garrigan et al. 2014), only lines *R31* and *R41* were *w*Mau-infected. We also screened the living copies of these lines for *wsp* using PCR, and both were infected, supporting reliable *w*Mau transmission in the lab since these lines were sampled in nature.

To assess the quality of our draft assemblies, we used BUSCO v. 3.0.0 to search for homologs of the near-universal, single-copy genes in the BUSCO proteobacteria database (Simao et al. 2015). As a control, we performed the same search using the reference genomes for *w*Ri (Klasson et al. 2009), *w*Au (Sutton et al. 2014), *w*Mel (Wu et al. 2004), *w*Ha (Ellegaard et al. 2013), and *w*No (Ellegaard et al. 2013).

### *Wolbachia* gene extraction and phylogenetics

To determine phylogenetic relationships and estimate divergence times, we obtained the public *Wolbachia* group-B genomes of: *w*AlbB that infects *Aedes albopictus* (Mavingui et al. 2012), *w*Pip_Pel that infects *Culex pipiens* (Klasson et al. 2008), *w*Pip_Mol that infects *Culex molestus* (Pinto et al. 2013), *w*No that infects *Drosophila simulans* (Ellegaard et al. 2013), and *w*VitB that infects *Nasonia vitripennis* (Kent et al. 2011); in addition to group-A genomes of: *w*Mel that infects *D. melanogaster* (Wu et al. 2004), *w*Suz that infects *D. suzukii* (Siozios et al. 2013), four *Wolbachia* that infect *Nomada* bees (*w*NFe, *w*NPa, *w*NLeu, and *w*NFa; Gerth and Bleidorn 2016), and three *Wolbachia* that infect *D. simulans* (*w*Ri, *w*Au and *w*Ha; Klasson *et al*. 2009; Sutton *et al*. 2014; Ellegaard *et al*. 2013). The previously published genomes and the five *w*Mau-infected *D. mauritiana* genomes were annotated with Prokka v. 1.11, which identifies homologs to known bacterial genes (Seemann 2014). To avoid pseudogenes and paralogs, we used only genes present in a single copy, and with no alignment gaps, in all of the genome sequences. Genes were identified as single copy if they uniquely matched a bacterial reference gene identified by Prokka v. 1.11. By requiring all homologs to have identical length in all of the draft *Wolbachia* genomes, we removed all loci with indels. 143 genes, a total of 113,943 bp, met these criteria when comparing all of these genomes. However, when our analysis was restricted to the five *w*Mau genomes, our criteria were met by 694 genes, totaling 704,613 bp. Including *w*No with the five *w*Mau genomes reduced our set to 671 genes with 682,494 bp. We calculated the percent differences for the three codon positions within *w*Mau and between *w*Mau and *w*No.

We estimated a Bayesian phylogram of the *Wolbachia* sequences with RevBayes 1.0.8 under the GTR + Γ model, partitioning by codon position (Höhna et al. 2016). Four independent runs were performed, which all agreed.

We estimated a chronogram from the *Wolbachia* sequences using the absolute chronogram procedure implemented in Turelli et al. (2018). Briefly, we generated a relative relaxed-clock chronogram with the GTR + Γ model with the root age fixed to 1 and the data partitioned by codon position. The relaxed clock branch rate prior was Γ(2,2). We used substitution-rate estimates of Γ(7,7) × 6.87×10^−9^ substitutions/3^rd^ position site/year to transform the relative chronogram into an absolute chronogram. This rate estimate was chosen so that the upper and lower credible intervals matched the posterior distribution estimated by Richardson et al. (2012), assuming 10 generations/year, normalized by their median estimate of 6.87×10^−9^ substitutions/3^rd^ position site/year. Although our relaxed-clock analyses allow for variation in substitution rates across branches, our conversion to absolute time depends on the unverified assumption that the median substitution rate estimated by Richardson et al. (2012) for *w*Mel is relevant across these broadly diverged *Wolbachia*. (To assess the robustness of our conclusions to model assumptions, we also performed a strict-clock analysis and a relaxed-clock analysis with branch-rate prior Γ(7,7).) For each analysis, four independent runs were performed, which all agreed. Our analyses all support *w*No as sister to *w*Mau.

We also estimated a relative chronogram for the host species using the procedure implemented in Turelli et al. (2018). Our host phylogeny was based on the same 20 nuclear genes used in Turelli et al. (2018): *aconitase, aldolase, bicoid, ebony, enolase, esc, g6pdh, glyp, glys, ninaE, pepck, pgi, pgm, pic, ptc, tpi, transaldolase, white, wingless* and *yellow*.

### Analysis of *Wolbachia* and mitochondrial genomes

We looked for copy number variation (CNV) between *w*Mau and its closest relative, *w*No across the whole *w*No genome. Reads from the five infected *w*Mau lines were aligned to the *w*No reference (Ellegaard et al. 2013) with bwa 0.7.12 (Li and Durbin 2009). We calculated the normalized read depth for each alignment over sliding 1,000-bp windows by dividing the average depth in the window by the average depth over the entire *w*No genome. The results were plotted and visually inspected for putative copy number variants (CNVs). The locations of CNVs were specifically identified with ControlFREEC v. 11.5 (Boeva et al. 2012), using a ploidy of one and a window size of 1,000. We calculated *P*-values for each identified CNV with the Wilcoxon Rank Sum and the Kolmogorov-Smirnov tests implemented in ControlFREEC.

We used BLAST to search for pairs of CI-factor (*cif*) homologs in *w*Mau and *w*No genomes that are associated with CI (Beckmann and Fallon 2013; Beckmann et al. 2017; LePage et al. 2017; Lindsey et al. 2018; Beckmann et al. 2019). (We adopt Beckmann et al. (2019)’s nomenclature that assigns names to loci based on their predicted enzymatic function, with superscripts denoting the focal *Wolbachia* strain.) These include predicted CI-inducing deubiquitylase (*cid*) *w*Pip_0282-*w*Pip_0283 (*cidA-cidB*^*w*Pip^) and CI-inducing nuclease (*cin*) wPip_0294-wPip_0295 (*cinA-cinB*^*w*Pip^) pairs that induce toxicity and rescue when expressed/co-expressed in *Saccharomyces cerevisiae* (Beckmann et al. 2017 and Beckmann et al. 2019); WD0631-WD632 (*cidA-cidB*^*w*Mel^) that recapitulate CI when transgenically expressed in *D. melanogaster* (LePage *et al*. 2017); and *w*No_RS01055 and *w*No_RS01050 that have been identified as a Type III *cifA-cifB* pair in the *w*No genome (LePage et al. 2017; Lindsey et al. 2018). *w*No_RS01055 and *w*No_RS01050 are highly diverged from *cidA-cidB*^*w*Mel^ and *cidA-cidB*^*w*Pip^ homologs and from *cinA-cinB*^*w*Pip^; however, this *w*No pair is more similar to *cinA-cinB*^*w*Pip^ in terms of protein domains, lacking a ubiquitin-like protease domain (Lindsey et al. 2018). We refer to these loci as *cinA-cinB*^*w*No^.

We found only homologs of the *cinA-cinB*^*w*No^ pair in *w*Mau genomes, which we extracted from our draft *w*Mau assemblies and aligned with MAFFT v. 7 (Katoh and Standley 2013). We compared *cinA-cinB*^*w*No^ to the *w*Mau homologs to identify single nucleotide variants (SNVs) among our *w*Mau assemblies.

*D. mauritiana* carry either the *ma*I mitochondrial haplotype, associated with *w*Mau infections, or the *ma*II haplotype (Rousset and Solignac 1995; Ballard 2000a; James and Ballard 2000). To determine the mitochondrial haplotype of each *D. mauritiana* line, we assembled the mitochondrial genomes by down-sampling the reads by a factor of 100, then assembling with ABySS 2.0.2 using a *K* value of 71 for our data (150 bp reads) and 35 for the published data (76 bp reads) (Garrigan et al. 2014). Down-sampling reads prevents the nuclear genome from assembling but does not inhibit assembly of the mitochondrial genome, which has much higher coverage. We deemed the mitochondrial assembly complete if all 13 protein-coding genes were present on the same contig and in the same order as in *D. melanogaster*. If the first attempt did not produce a complete mitochondrial assembly, we adjusted the down-sampling fraction until a complete assembly was produced for each line.

Annotated reference mitochondrial sequences for the *D. mauritiana* mitochondrial haplotypes *ma*I and *ma*II were obtained from Ballard et al. (2000b), and the 13 protein-coding genes were extracted from our assemblies using BLAST and aligned to these references. The *ma*I and *ma*II reference sequences differ at 343 nucleotides over these protein-coding regions. We identified our lines as carrying the *ma*I haplotype if they differed by fewer than five nucleotides from the *ma*I reference and as *ma*II if they differed by fewer than five nucleotides from the *ma*II reference. None of our assemblies differed from both references at five or more nucleotides.

### *w*Mau phenotypic analyses

Previous analyses have demonstrated that *w*Mau does not cause CI (Giordano et al. 1995). To check the generality of this result, we reciprocally crossed *w*Mau-infected *R31 D. mauritiana* with uninfected *R4* and measured egg hatch. Flies were reared under controlled conditions at 25°C for multiple generations leading up to the experiment. We paired 1–2-day-old virgin females with 1–2-day-old males in a vial containing spoons with cornmeal media and yeast paste. After 24 hr, pairs were transferred to new spoons, and this process was repeated for five days. Eggs on each spoon were given 24 hr at 25°C to hatch after flies were removed. To test for CI, we used nonparametric Wilcoxon tests to compare egg hatch between reciprocal crosses that produced at least 10 eggs. All experiments were carried out at 25°C with a 12 light:12 dark photoperiod.

To determine if *w*Mau generally enhances *D. mauritiana* fecundity, we assessed the fecundity of two *w*Mau-infected isofemale lines from Mauritius (*R31* and *R41*); we also reciprocally introgressed *w*Mau from each of these lines to assess host effects. To do this we crossed *R31* females with *R41* males and backcrossed F_1_ females to *R41* males—this was repeated for four generations to generate the reciprocally introgressed *R41*^*R31*^ genotypes (*w*Mau variant denoted by superscripts). A similar approach was taken to generate *R31*^*R41*^ genotypes. This approach has previously revealed *D. teissieri*-host effects on *w*Tei-induced CI (Cooper et al. 2017). To assay fecundity, we reciprocally crossed each genotype (*R31, R41, R31*^*R41*^, *R31*^*R41*^ ) to uninfected line *R4* to generate paired infected- and uninfected-F_1_ females with similar genetic backgrounds. The *w*Mau-infected and uninfected F_1_ females were collected as virgins and placed in holding vials. We paired 3–7-day-old females individually with an uninfected-*R4* male (to stimulate oviposition) in vials containing a small spoon filled with standard cornmeal medium and coated with a thin layer of yeast paste. We allowed females to lay eggs for 24 hours, after which pairs were transferred to new vials. This was repeated for five days. At the end of each 24-hr period, spoons were frozen until counted. All experiments were carried out at 25°C with a 12 light:12 dark photoperiod.

We also measured egg lay of *w*Mau-infected (*R31*) and tetracycline-cleared uninfected (*R31-tet*) genotypes over 24 days, on apple-agar plates, to more closely mimic the methods of Fast et al. (2011). We fed flies 0.03% tetracycline concentrated medium for four generations to generate the *R31-tet* genotype. We screened F_1_ and F_2_ individuals for *w*Mau using PCR, and we then fed flies tetracycline food for two additional generations. In the fourth generation, we assessed *w*Mau titer using qPCR to confirm that each genotype was cleared of *w*Mau infection. We reconstituted the gut microbiome by rearing *R31-tet* flies on food where *R31* males had fed and defecated for 48 hours. Flies were given at least three more generations to avoid detrimental effects of tetracycline treatment on mitochondrial function (Ballard and Melvin 2007). We then paired individual 6–7-day-old virgin *R31* (*N* = 30) and *R31-tet* (*N* = 30) females in bottles on yeasted apple-juice agar plates with an *R4* male to stimulate oviposition. Pairs were placed on new egg-lay plates every 24 hrs. After two weeks, we added one or two additional *R4* males to each bottle to replace any dead males and to ensure that females were not sperm limited as they aged.

We used nonparametric Wilcoxon tests to assess *w*Mau effects on host fecundity. We then estimated the fitness parameter *F* in the standard discrete-generation model of CI (Hoffmann et al. 1990; Turelli 1994). We used the ‘*pwr.t2n.test*’ function in the ‘*pwr*’ library in R to assess the power of our data to detect increases to *F*. Pairs that laid fewer than 10 eggs across each experiment were excluded from analyses, but our results are robust to this threshold.

To estimate the fidelity of maternal transmission, *R31* and *R41* females were reared at 25°C for several generations prior to our experiment. In the experimental generation, 3-5 day old inseminated females were placed individually in vials that also contained two males. These *R31* (*N* = 17) and *R41* (*N* = 19) sublines were allowed to lay eggs for one week. In the following generation we screened F1 offspring for *w*Mau infection using PCR as described above.

## RESULTS

### *Wolbachia* infection status

Out of 32 *D. mauritiana* lines that we analyzed, 11 were infected with *w*Mau *Wolbachia* (infection frequency = 0.34; binomial confidence interval: 0.19, 0.53). In contrast, none of the *D. simulans* stocks (*N* = 4) sampled from Madagascar were infected, precluding our ability to directly compare *w*Mau and *w*Ma. Our new *w*Mau frequency estimate is not statistically different from a previous estimate (Giordano et al. 1995: infection frequency, 0.46; binomial confidence interval, (0.34, 0.58); Fisher’s Exact Test, *P* = 0.293), based largely on assaying a heterogenous collecton of stocks in various laboratories. These relatively low infection frequencies are consistent with theoretical expectations given that *w*Mau does not cause CI (Giordano et al. 1995; our data reported below). The intermediate *w*Mau frequencies on Mauritius suggest that *w*Mau persists at a balance between positive effects on host fitness and imperfect maternal transmission. Quantitative predictions, based on the idealized model of Hoffmann and Turelli (1997), are discussed below. The maintenance of *w*Mau is potentially analogous to the persistence of other non-CI-causing *Wolbachia*, specifically *w*Au in some Australian populations of *D. simulans* (Hoffmann et al. 1996; Kriesner et al. 2013) and *w*Suz in *D. suzukii* and *w*Spc in *D. subpulchrella* (Hamm et al. 2014; Conner et al. 2017; Turelli et al. 2018; but see Cattel et al. 2018).

### Draft *w*Mau genome assemblies and comparison to *w*No

The five *w*Mau draft genomes we assembled were of very similar quality (Supplemental Table 1). N50 values ranged from 60,027 to 63,676 base pairs, and our assemblies varied in total length from 1,266,004 bases to 1,303,156 bases (Supplemental Table 1). Our BUSCO search found exactly the same genes in each draft assembly, and the presence/absence of genes in our *w*Mau assemblies was comparable to those in the complete genomes used as controls (Supplemental Table 2). In comparing our five *w*Mau draft genomes over 694 single-copy, equal-length loci comprising 704,613 bp, we found only one SNP. Four sequences (*R9, R31, R41* and *R60*) are identical at all 704,613 bp. *R29* differs from them at a single nucleotide, a nonsynonymous substitution in a locus which Prokka v. 1.11 annotates as “bifunctional DNA-directed RNA polymerase subunit beta/beta.”

**Table 1.**
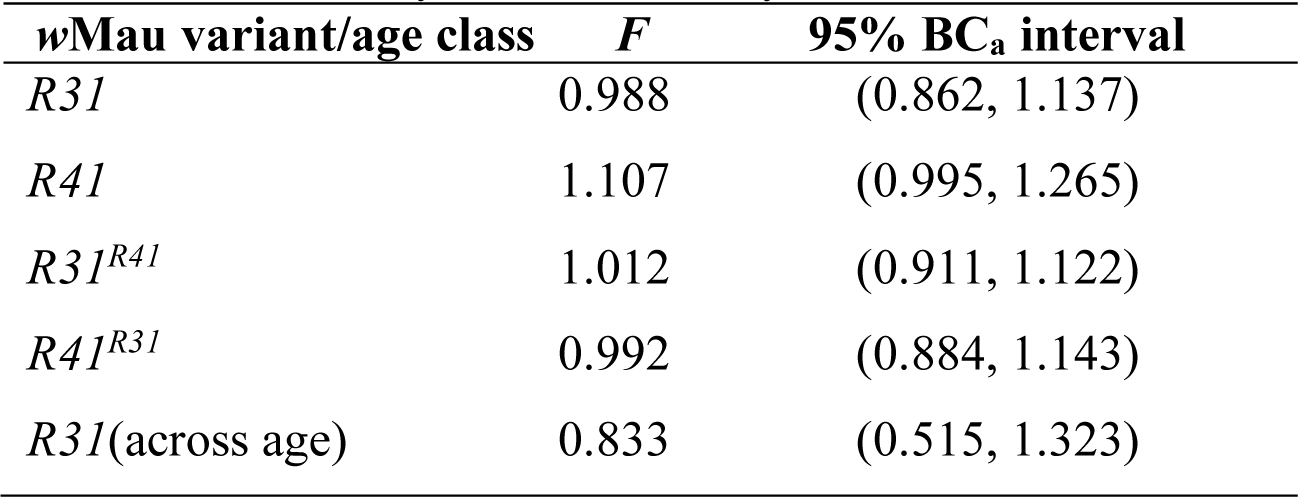
Estimates of the relative fitness parameter *F* indicate that *w*Mau fecundity effects are likely to be minimal.

Comparing these five *w*Mau sequences to the *w*No reference (Ellegaard et al. 2013) over 671 genes with 682,494 bp, they differ by 0.068% overall, with equivalent divergence at all three codon positions (0.067%, 0.061%, and 0.076%, respectively).

### *Wolbachia* phylogenetics

As expected from the sequence comparisons, our group-B phylogram places *w*Mau sister to *w*No (Figure 1A). This is consistent with previous analyses using fewer loci that placed *w*Mau (or *w*Ma in *D. simulans*) sister to *w*No (James and Ballard 2000; Zabalou et al. 2008; Toomey et al. 2013). Our chronogram (Figure 1B) estimates the 95% credible interval for the split between the group-B versus group-A *Wolbachia* strains as 6 to 36 mya (point estimate, 16 mya). Reducing the variance on the substitution-rate-variation prior by using Γ(7,7) rather than Γ(2,2), changes the credible interval for the A-B split to 8 to 46 mya (point estimate, 21 mya). In contrast, a strict clock analysis produces a credible interval of 12 to 64 mya (point estimate, 31 mya). These estimates are roughly comparable to an earlier result based on a general approximation for the synonymous substitution rate in bacteria (Ochman and Wilson 1987) and data from only the *ftsZ* locus (59–67 mya, Werren et al. 1995). However, our estimates are much lower than an alternative estimate based on comparative genomics (217 mya, Gerth and Bleidorn 2016). We discuss this discrepancy below.

**Figure 1.**
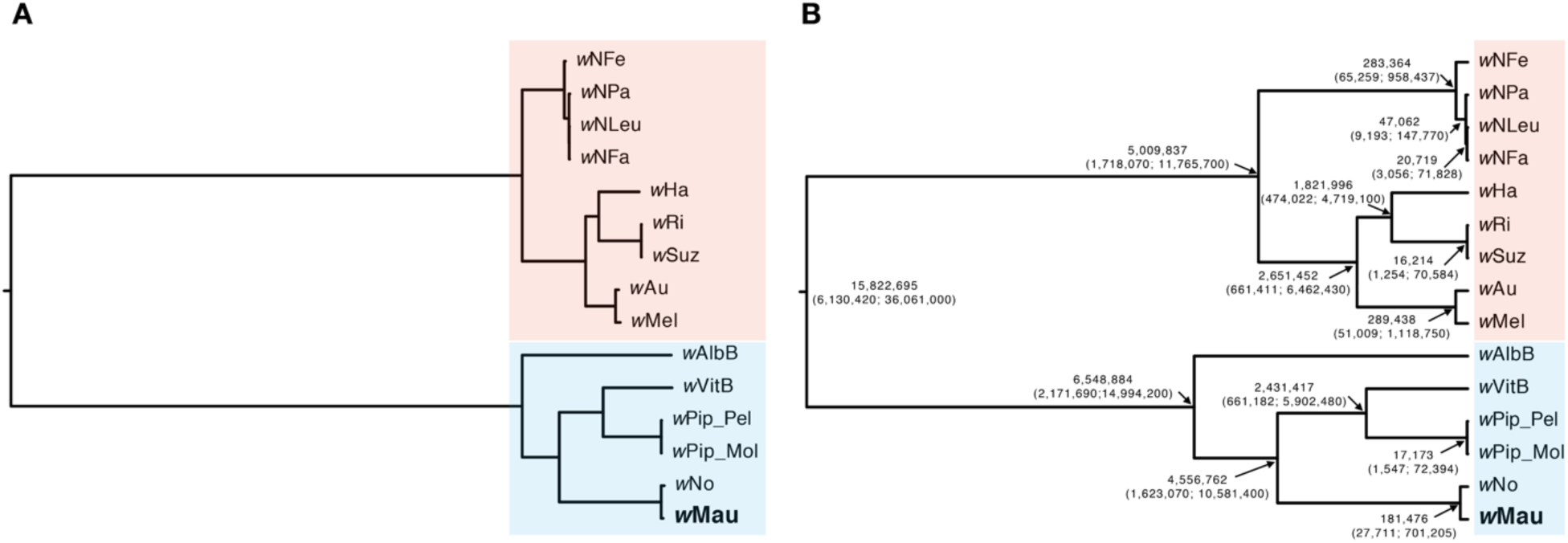
A) An estimated phylogram for various group-A (red) and group-B (blue) *Wolbachia* strains. All nodes have Bayesian posterior probabilities of 1. The phylogram shows significant variation in the substitution rates across branches, with long branches separating the A and B clades. **B)** An estimated chronogram for the same strains, with estimated divergence times and their confidence intervals at each node. To obtain these estimates, we generated a relative relaxed-clock chronogram with the GTR + Γ model with the root age fixed to 1, the data partitioned by codon position, and with a Γ(2,2) branch rate prior. We used substitution-rate estimates of Γ(7,7) × 6.87×10^−9^ substitutions/3^rd^ position site/year to transform the relative chronogram into an absolute chronogram.

The observed divergence between *w*No and *w*Mau is consistent across all three codon positions, similar to other recent *Wolbachia* splits like that between *w*Ri and *w*Suz (Turelli et al. 2018). Conversely, observed divergence at each codon position generally varies across the chronogram, leading to inflation of the *w*No-*w*Mau (181,476 years; credible interval = 27,711 to 701,205 years; Figure 1B) and *w*Ri-*w*Suz (16,214; credible interval = 1,254 to 70,584) divergence point estimates; the latter is about 1.6 times as large as the value in Turelli et al. (2018). (Nevertheless, the confidence intervals of our and Turelli et al. (2018)’s *w*Ri-*w*Suz divergence estimates overlap.) To obtain an alternative estimate of *w*No-*w*Mau divergence, we estimated divergence time using the observed third-position pairwise divergence (0.077%, or 0.039% from tip to MRCA) and Richardson et al. (2012)’s estimate of the “short-term evolutionary rate” of *Wolbachia* third-position divergence within *w*Mel. This approach produces a point estimate of 57,000 years, with a credible interval of 30,000 to 135,000 years for the *w*No-*w*Mau split. In Cooper et al. (2019), we address in detail how a constant substitution-rate ratio among codon positions across the tree, assumed by the model, affects these estimates.

### Analysis of *Wolbachia* and mitochondrial genomes

We looked for CNVs in *w*Mau relative to sister *w*No by plotting normalized read depth along the *w*No genome. There were no differences in copy number among the *w*Mau variants, but compared to *w*No, ControlFREEC identified four regions deleted from all *w*Mau that were significant according to the Wilcoxon Rank Sum and Kolmogorov-Smirnov tests (Figure 2 and Supplemental Table 3). These deleted regions of the *w*Mau genomes include many genes, including many phage-related loci, providing interesting candidates for future work (listed in Supplementary Table 4).

**Figure 2.**
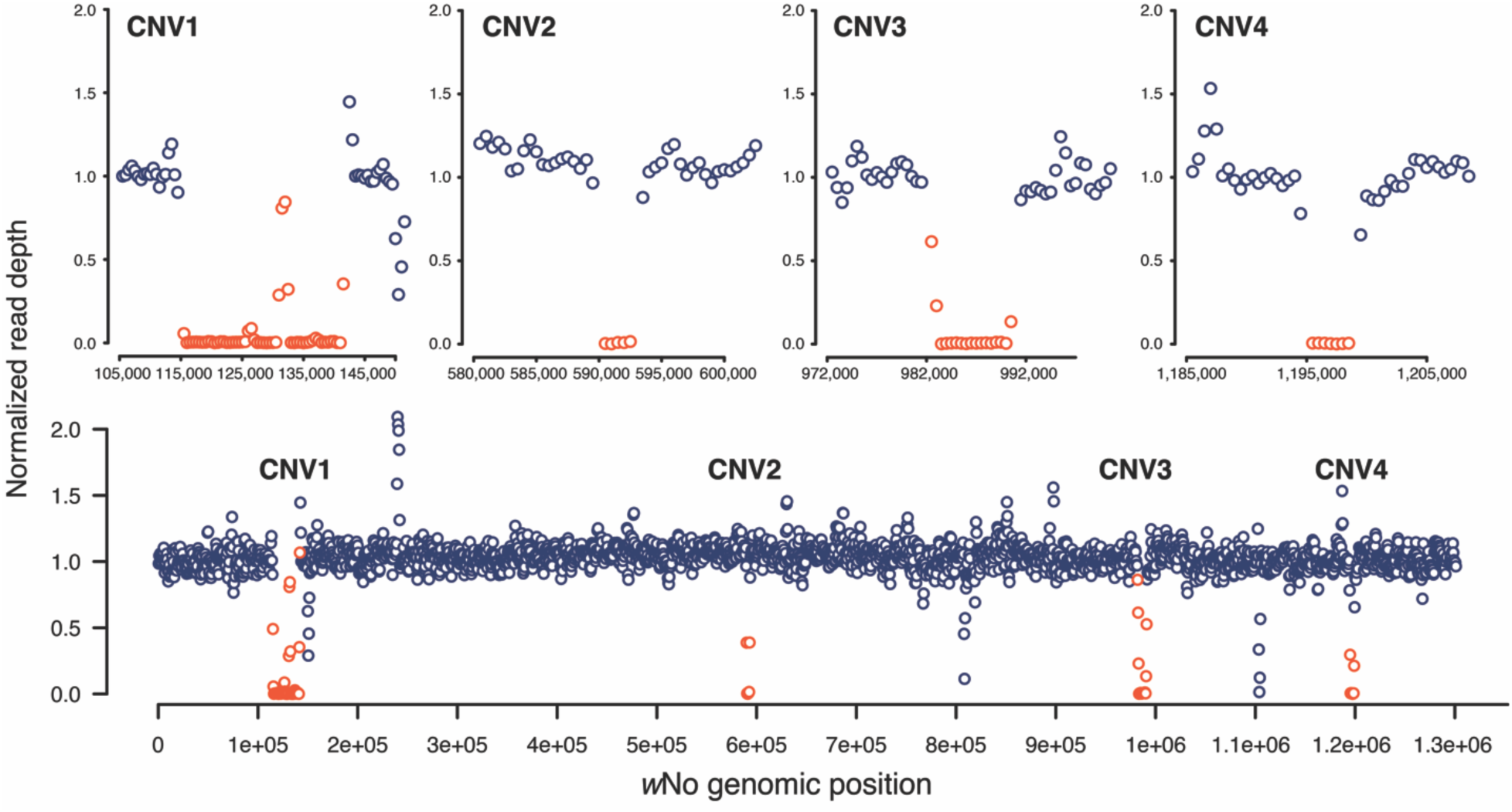
All *w*Mau variants share four large deletions, relative to sister *w*No. Top panel) The normalized read depth for *w*Mau *R60* plotted across the four focal regions of the *w*No reference genome; 10 kb of sequence surrounding regions are plotted on either side of each region. Bottom panel) The normalized read depth of *w*Mau *R60* plotted across the whole *w*No reference genome. Regions that do not contain statistically significant CNVs are plotted in dark blue, and regions with significant CNVs are plotted in red. All *w*Mau variants share the same CNVs, relative to *w*No.

To test the hypothesis that *cif* loci are disrupted, we searched for pairs of loci known to be associated with CI and found homologs to the *cinA-cinB*^*w*No^ pair in each of our draft assemblies, but we did not find homologs to the *cidA-cidB*^*w*Mel^, *cidA-cidB*^*w*Pip^, or to the *cinA-cinB*^*w*Pip^ pairs. There were no variable sites in *cinA-cinB*^*w*No^ homologs among our five *w*Mau assemblies. Relative to *w*No, all *w*Mau variants share a one base pair deletion at base 1133 out of 2091 (amino acid 378) in the *cinB*^*w*No^ homolog. This frameshift introduces over 10 stop codons, with the first at amino acid 388, potentially making this predicted CI-causing-toxin protein nonfunctional. We also identified a nonsynonymous substitution in amino acid 264 of the *cinB*^*w*No^ homolog (*w*No codon ACA, Thr; *w*Mau codon AUA, Ile) and two SNVs in the region homologous to *cinA*^*w*No^ : a synonymous substitution in amino acid 365 (*w*No codon GUC, *w*Mau codon GUU) and a nonsynonymous substitution in amino acid 397 (*w*No codon GCU, amino acid Ala; *w*Mau codon GAU, amino acid Asp). Disruption of CI is consistent with theoretical analyses showing that selection within a host species does not act directly on the level of CI (Prout 1994; Turelli 1994; Haygood and Turelli 2009). Future functional analyses will determine whether disruption of regions homologous to *cinA-cinB*^*w*No^ underlie the lack of *w*Mau CI.

Of the *D. mauritiana* lines tested (*N* = 9), one line (uninfected-*R44*) carries the *ma*II mitochondrial haplotype, while the other eight carry *ma*I. Rousset and Solignac (1995) reported a similar *ma*II frequency, with 3 of 26 lines sampled in 1985 carrying *ma*II. The *ma*I and *ma*II references differ by 343 SNVs across the proteome, and *R44* differs from the *ma*II reference by 4 SNVs in the proteome. Four of our *ma*I lines (*R23, R29, R32*, and *R39*) are identical to the *ma*I reference, while three (*R31, R41*, and *R60*) have one SNV and one (*R9*) has two SNVs relative to *ma*I reference. One SNV is shared between *R9* and *R60*, but the other three SNVs are unique. Our results agree with past analyses that found *w*Mau is perfectly associated with the *ma*I mitochondrial haplotype (Rousset and Solignac 1995; Ballard 2000a; James and Ballard 2000). The presence of *ma*II among the uninfected is interesting. In contrast to *ma*I, which is associated with introgression with *D. simulans* (Ballard 2000a; James and Ballard 2000), *ma*II appears as an outgroup on the mtDNA phylogeny of the *D. simulans* clade and is not associated with *Wolbachia* (Ballard 2000b, Fig. 5; James and Ballard 2000). Whether or not *Wolbachia* cause CI, if they are maintained by selection-imperfect-transmission balance, we expect all uninfected flies to eventually carry the mtDNA associated with infected mothers (Turelli et al. 1992). We present a mathematical analysis of the persistence of *ma*II below.

### Analysis of *w*Mau phenotypes

In agreement with Giordano et al. (1995), we found no difference between the egg hatch of uninfected females crossed to *w*Mau-infected males (0.34 ± 0.23 SD, *N* = 25) and the reciprocal cross (0.29 ± 0.28 SD, *N* = 24), indicating no CI. In contrast to Fast et al. (2011), we find no evidence that *w*Mau affects *D. mauritiana* fecundity (Supplemental Table 5 and Figure 3), regardless of host genetic backgrounds. Across both experiments assessing *w*Mau fecundity effects in their natural backgrounds (*R31* and *R41*), we counted 27,221 eggs and found no difference in the number of eggs laid by infected (mean = 238.20, SD = 52.35, *N* = 60) versus uninfected (mean = 226.82, SD = 67.21, *N* = 57) females over the five days of egg lay (Wilcoxon test, *W* = 1540.5, *P* = 0.357); and across both experiments that assessed *w*Mau fecundity effects in novel host backgrounds (*R31*^*R41*^ and *R41*^*R31*^), we counted 30,358 eggs and found no difference in the number of eggs laid by infected (mean = 253.30, SD = 51.99, *N* = 60) versus uninfected (mean = 252.67, SD = 63.53, *N* = 60) females over five days (Wilcoxon test, *W* = 1869.5, *P* = 0.719). [The mean number of eggs laid over five days, standard deviation (SD), sample size (*N*), and *P*-values from Wilcoxon tests are presented in Supplemental Table 5 for all pairs.]

**Figure 3.**
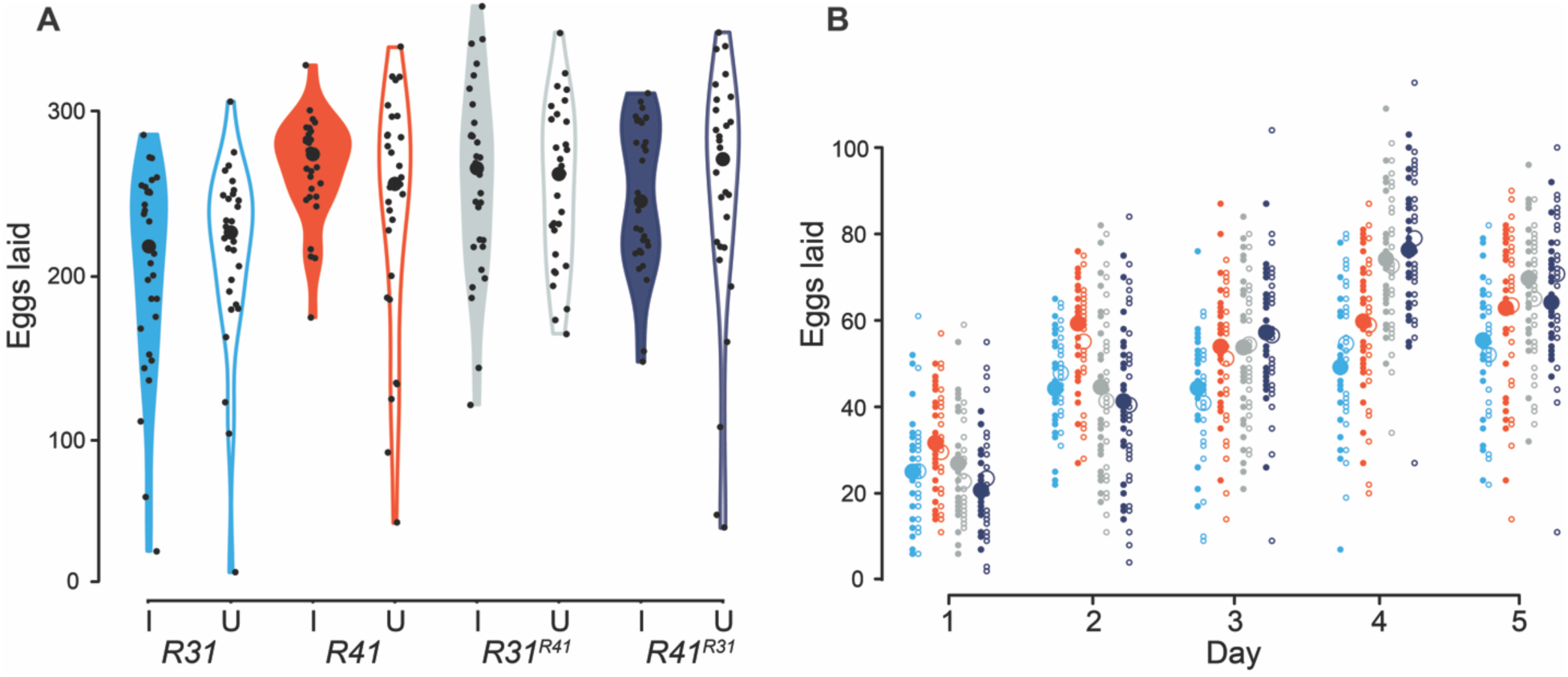
*w*Mau infections do not influence *D. mauritiana* fecundity, regardless of host genomic background. **A)** Violin plots of the number of eggs laid by *D. mauritiana* females over five days when infected with their natural *w*Mau variant (*R31I* and *R41I*), when infected with a novel *w*Mau variant (*R31*^*R41*^*I* and *R41*^*R31*^*I*), and when uninfected (*R31U, R41U, R31*^*R41*^*U*, and *R41*^*R31*^*U*). Large black dots are medians, and small black dots are the eggs laid by each replicate over five days. **B)** The daily egg lay of these same infected (solid circles) and uninfected (open circles) *R31* (aqua), *R41* (red), *R31*^*R41*^ (gray), and *R41*^*R31*^ (dark blue) genotypes is reported. Large circles are means of all replicates, and small circles are the raw data. Only days where females laid at least one egg are plotted. Cytoplasm sources are denoted by superscripts for the reciprocally introgressed strains.

We sought to determine if *w*Mau fecundity effects depend on host age with a separate experiment that assessed egg lay over 24 days on apple-agar plates, similar to Fast et al. (2011). Across all ages, we counted 9,459 eggs and found no difference in the number of eggs laid by infected (mean = 156.29, SD = 138.04, *N* = 28) versus uninfected (mean = 187.70, SD = 168.28, *N* = 27) females (Wilcoxon test, *W* = 409, *P* = 0.608) (Figure 4). While our point estimates indicate that *w*Mau does not increase host fecundity, egg lay was generally lower and more variable on agar plates relative to our analyses of egg lay on spoons described above.

**Figure 4.**
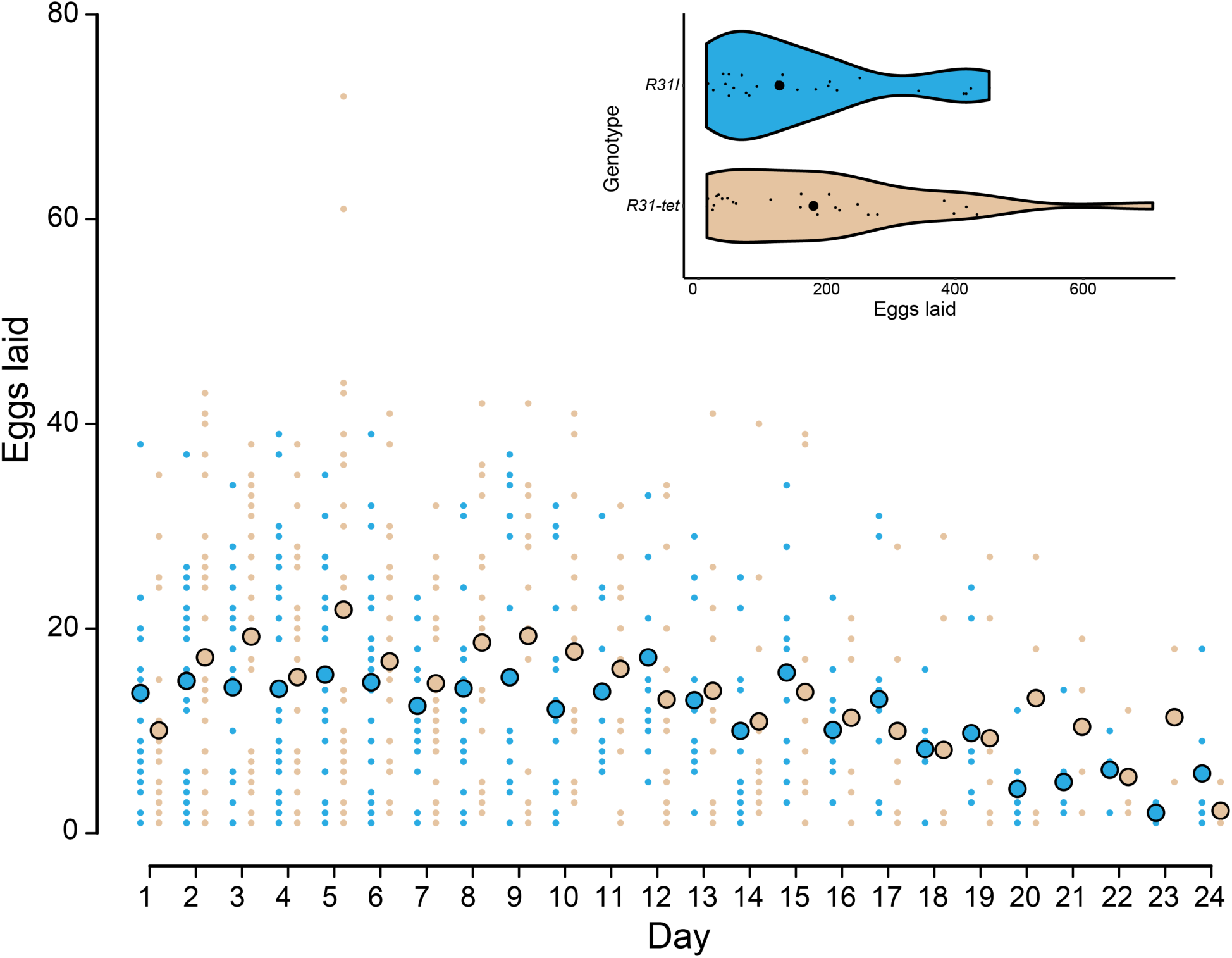
The mean number of eggs laid by infected *R31* (*R31I*, large aqua dots) and uninfected *R31-tet* (large tan dots) genotypes are similar. Egg counts for each replicate are also plotted (small dots). Violin plots show egg lay across all ages for each genotype; large black circles are medians, and small black circles are the number of eggs laid by each replicate.

**Figure 5.**
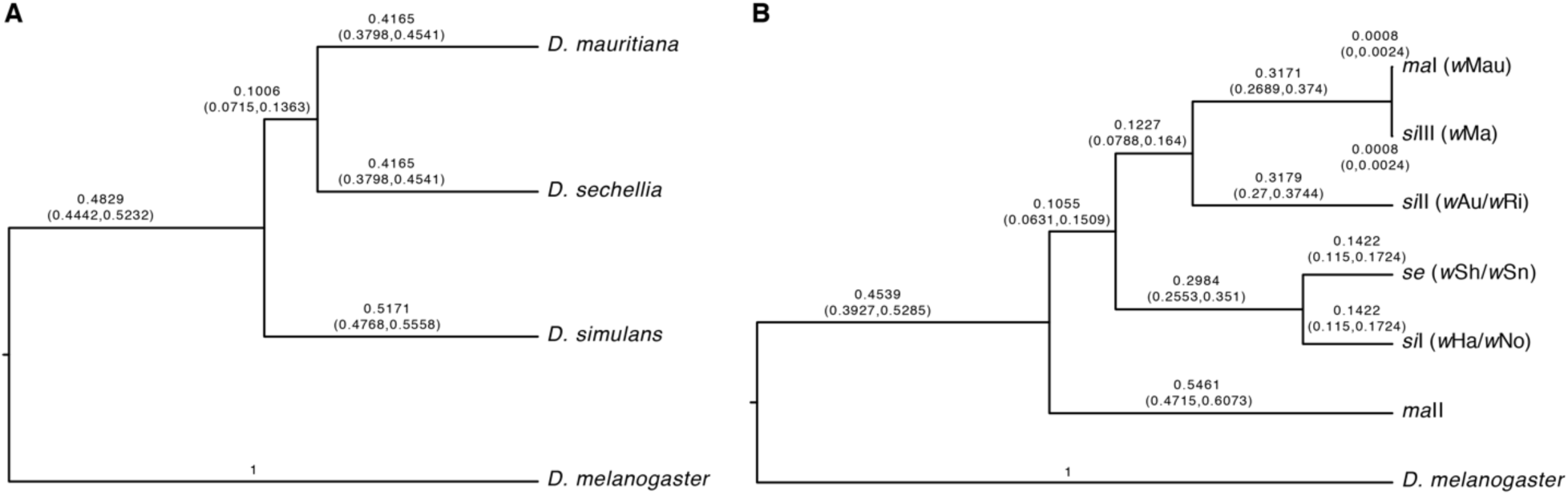
**A)** A nuclear relative chronogram. **B)** A mitochondrial relative chronogram with co-occurring *Wolbachia* strains listed in parentheses. See the text for an interpretation of the results, including the artifactual resolution of the phylogeny of the *D. simulans* clade.

With these data, we estimated the fitness parameter *F* in the standard discrete-generation model of CI (Hoffmann et al. 1990; Turelli 1994). Taking the ratio of replicate mean fecundity observed for *w*Mau-infected females to the replicate mean fecundity of uninfected females in naturally sampled *R31* and *R41 D. mauritiana* backgrounds, we estimated *F* = 1.05 (95% BC_a_ interval: 0.96, 1.16). Following reciprocal introgression of *w*Mau and host backgrounds (i.e., the *R31*^*R41*^ and *R41*^*R31*^ genotypes), we estimated *F* = 1.0 (95% BC_a_ interval: 0.93, 1.09). Finally, across all 24 days of our age-effects experiment, we estimated *F* = 0.83, 95% BC_a_ interval: 0.52, 1.32) for *R31*, which overlaps with our estimate of *F* for *R31* in our initial experiment (Table 1). BC_a_ confidence intervals were calculated using the two-sample acceleration constant given by equation 15.36 of Efron and Tibshirani (1993). (Estimates of *F* and the associated BC_a_ confidence intervals are reported in Table 1 for each genotype and condition.) Consistent with our other analyses, we find little evidence that *w*Mau significantly increases fecundity. However, our data do not have much statistical power to detect values of *F* on the order of 1.05, which may suffice to produce *F*(1 – *μ*) > 1 and deterministic spread of *w*Mau from low frequencies. We present our power calculations in Figure 1B of the Supplementary Information.

Finally, we assessed the fidelity of *w*Mau maternal transmission under standard laboratory conditions. We excluded sublines that produced fewer than 8 F1 offspring. In all cases, *R31* (*N* = 17) sublines produced offspring that were all infected, indicating perfect maternal transmission. In contrast, one *R41* subline produced one uninfected individual out of a total of 18 F1 offspring produced; all other *R41* sublines meeting our criteria (*N* = 15) produced only infected F1 offspring, resulting in nearly perfect maternal transmission across all *R41* sublines (*μ* = 0.0039).

### Mathematical analyses of *Wolbachia* frequencies and mtDNA polymorphism

If *Wolbachia* do not cause CI (or any other reproductive manipulation), their dynamics can be approximated by a discrete-generation haploid selection model. Following Hoffmann and Turelli (1997), we assume that the relative fecundity of *Wolbachia*-infected females is *F*, but a fraction *μ* of their ova do not carry *Wolbachia*. Given our ignorance of the nature of *Wolbachia*’s favorable effects, the *F* represents an approximation for all fitness benefits. If *F*(1 – *μ*) > 1, the equilibrium *Wolbachia* frequency among adults is

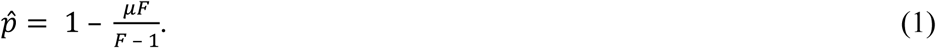

Imperfect maternal transmisson has been documented for field-collected *D. simulans* infected with *w*Ri (Hoffmann and Turelli 1988; Turelli and Hoffmann 1995; Carrington et al. 2011), *D. melanogaster* infected with *w*Mel (Hoffmann et al. 1998) and *D. suzukii* infected with *w*Suz (Hamm et al. 2014). The estimates range from about 0.01 to 0.1. Given that we have documented imperfect maternal transmission of *w*Mau in the laboratory, we expect (more) imperfect transmission in nature (Turelli and Hoffmann 1995; Carrington et al. 2011). In order for the equilibrium *Wolbachia* frequency to be below 0.5, approximation (1) requires that the relative fecundity of infected females satisfies

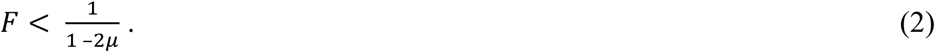

Thus, even for *μ* as large a 0.15, which greatly exceeds our laboratory estimates for *w*Mau and essentially all estimates of maternal transmission failure from nature, *Wolbachia* can increase fitness by at most 43% and produce an equilibrium frequency below 0.5 (Supplemental Figure 1A). Conversely, (1) implies that a doubling of relative fecundity by *Wolbachia* would produce an equilibrium frequency 1 – 2*μ*. If *μ* ≤ 0.25, consistent with all available data, the predicted equilibrium implied by a *Wolbachia*-induced fitness doubling significantly exceeds the observed frequency of *w*Mau. Hence, a four-fold fecundity effect, as described by Fast et al. (2011), is inconsistent with the frequency of *w*Mau in natural populations of *D. mauritiana*. Field estimates of *μ* for *D. mauritiana* will provide better theory-based bounds on *w*Mau fitness effects that would be consistent with *w*Mau tending to increase when rare on Mauritius, i.e., conditions for *F*(1 – *μ*) > 1.

Our theoretical analysis, addressing the plausibility of a four-fold fitness increase caused by *w*Mau, assumes that the observed frequency of *w*Mau approximates selection-transmission equilibrium, as described by (1). With only two frequency estimates (one from a heterogeneous collection of laboratory stocks), we do not know that the current low frequency is temporarlly stable. Also, we do not know that the mutations we detect in *cinA-cinB*^*w*No^ homologs are responsible for the lack of *w*Mau CI. One alternative is that *D. mauritiana* has evolved to suppress CI (for host suppression of male killing, see Hornet et al. 2006 and Vanthournout and Hendrickx 2016). Host suppression of CI is expected (Turelli 1994), and it may explain the low CI caused by *w*Mel in *D. melanogaster* (Hoffmann and Turelli 1997). However, the fact that *w*Mau does not produce CI in *D. simulans*, a host that allows *w*Mel and other strains to induce strong CI even though little CI is produced in their native hosts, argues against host suppression as the explanation for the lack of CI caused by *w*Mau in *D. mauritiana*. Nevertheless, the loss of CI from *w*Mau may be quite recent; and *w*Mau may be on its way to elimination in *D. mauritiana*. If so, our equilibrium analysis is irrelevant – but this gradual-loss scenario is equally inconsistent with the four-fold fecundity effect proposed by Fast et al. (2011).

As noted by Turelli et al. (1992), if *Wolbachia* is introduced into a population along with a diagnostic mtDNA haplotype that has no effect on fitness, imperfect *Wolbachia* maternal transmission implies that all infected and uninfected individuals will eventually carry the *Wolbachia*-associated mtDNA, because all will have had *Wolbachia*-infected maternal ancestors. We conjectured that a stable mtDNA polymorphism might be maintained if *Wolbachia*-associated mtDNA introduced by introgression is deleterious in its new nuclear background. We refute our conjecture in Appendix 1. We show that the condition for *Wolbachia* to increase when rare, *F*(1 – *μ*) > 1, ensures that the native mtDNA will be completely displaced by the *Wolbachia*-associated mtDNA, even if it lowers host fitness once separated from *Wolbachia*.

How fast is the mtDNA turnover, among *Wolbachia*-uninfected individuals, as a new *Wolbachia* invades? This is easiest to analyze when the mtDNA introduced with *Wolbachia* has no effect on fitness, so that the relative fitness of *Wolbachia*-infected versus uninfected individuals is *F*, irrespective of the mtDNA haplotype of the uninfected individuals. As shown in Appendix 1, the frequency of the ancestral mtDNA haplotype among uninfected individuals, denoted *r*_t_, declines as

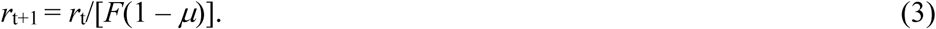

Assuming *r*_0_ = 1, recursion (3) implies that even if *F*(1 – *μ*) is only 1.01, the frequency of the ancestral mtDNA haplotype should fall below 10^−4^ after 1000 generations. A much more rapid mtDNA turnover was seen as the CI-causing *w*Ri swept northward through California populations of *D. simulans* (Turelli et al. 1992; Turelli and Hoffmann 1995). Thus, it is theoretically unexpected, under this simple model, that mtDNA haplotype *ma*II, which seems to be ancestral in *D. mauritiana* (Rousset and Solignac 1995; Ballard 2000a), persists among *Wolbachia*-uninfected *D. mauritiana*, given that all sampled *Wolbachia*-infected individuals carry *ma*I. However, spatial variation in fitnesses is one possible explanation for this polymorphism (Gliddon and Strobeck 1975), which has persisted since at least 1985.

## DISCUSSION

### *w*Mau is sister to *w*No and diverged from group-A *Wolbachia* less than 100 mya

Our phylogenetic analyses place *w*Mau sister to *w*No, in agreement with past analyses using fewer data (James and Ballard 2000; Zabalou et al. 2008; Toomey et al. 2013). The relationships we infer agree with those from recently published phylograms (Gerth and Bleidorn 2016; Lindsey et al. 2018) (Figure 1A).

Depending on the prior used for substitution-rate variation, we estimate that *w*Mau and other group-B *Wolbachia* diverged from group-A strains about 6–46 mya. This is roughly consistent with a prior estimate using only *ftsZ* (58–67 mya, Werren et al. 1995), but is inconsistent with a recent estimate using 179,763 bases across 252 loci (76–460 mya, Gerth and Bleidorn 2016). There are several reasons why we question the Gerth and Bleidorn (2016) calibration. First, Gerth and Bleidorn (2016)’s chronogram placed *w*No sister to all other group-B *Wolbachia*, in disagreement with their own phylogram (Gerth and Bleidorn 2016, Figure 3). In contrast, our phylogram and that of Lindsey et al. (2018) support *w*AlbB splitting from all other strains at this node. Second, the Gerth and Bleidorn (2016) calibration estimated the split between *w*Ri that infects *D. simulans* and *w*Suz that infects *D. suzukii* at 900,000 years. This estimate is more than an order of magnitude higher than ours (16,214 years) and nearly two orders of magnitude higher than the 11,000 year estimate of Turelli et al. (2018) who found 0.014% third position divergence between *w*Ri and *w*Suz (i.e., 0.007% along each branch) over 506,307 bases. Raychoudhury et al. (2009) and Richardson et al. (2012) both estimated a rate of about 7 × 10^−9^ substitutions/3^rd^ position site/year between *Wolbachia* in *Nasonia* wasps and within *w*Mel, respectively. An estimate of 900,000 years requires a rate about 100 times slower, 7.8 × 10^−11^ substitutions/3^rd^ position site/year, which seems implausible. Finally, using data kindly provided by Michael Gerth, additional analyses indicate that the third-position rates required for the *Wolbachia* divergence times estimated by Gerth and Bleidorn (2016) between *Nomada flava* and *N. leucophthalma* (1.72 × 10^−10^), *N. flava* and *N. panzeri* (3.78 × 10^−10^) (their calibration point), and *N. flava* and *N. ferruginata* (4.14 × 10^−10^) are each more than 10 times slower than those estimated by Raychoudhury et al. (2009) and Richardson et al. (2012), which seems unlikely. Our analyses suggest that the A-B group split occurred less than 100 mya.

### The lack of CI is consistent with intermediate *w*Mau infection frequencies

Across 671 genes (682,494 bases), the *w*Mau genomes were identical and differed from *w*No by only 0.068%. Across the coding regions we analyzed, we found few SNVs and no CNVs among *w*Mau variants. Our analyses did identify four large deletions shared by all *w*Mau genomes, relative to *w*No. Despite the close relationship between *w*Mau and *w*No, *w*No causes CI while *w*Mau does not (Giordano et al. 1995; Merçot et al. 1995; Rousset and Solignac 1995, our data). We searched for all pairs of loci known to cause CI and found only homologs to the *cinA-cinB*^*w*No^ pair in *w*Mau genomes. All *w*Mau variants share a one-base-pair deletion in the *w*Mau region homologous to *cinB*^*w*No^. This mutation introduces a frameshift and more than ten stop codons. Future functional work will help determine if disruption of this predicted-toxin locus underlies the lack of CI in *w*Mau. Regardless, the lack of CI is consistent with the prediction that selection within host lineages does not directly act on the intensity of CI (Prout 1994; Turelli 1994). We predict that analysis of additional non-CI-causing strains will reveal additional examples of genomic remnants of CI loci. Among non-CI *Wolbachia*, the relative frequency of those with non-functional CI loci, versus no CI loci, is unknown.

Irrespective of whether CI was lost or never gained, non-CI *Wolbachia* have lower expected equilibrium infection frequencies than do CI-causing variants (Kriesner et al. 2016). The *w*Mau infection frequency of approximately 0.34 on Mauritius (Giordano et al. 1995; our data) is consistent with this prediction. Additional sampling of Mauritius, preferably over decades, will determine whether intermediate *w*Mau frequencies are temporally stable. Such temporal stability depends greatly on values of *F* and *μ* through time suggesting additional field-based estimates of these parameters will be useful.

*w*Mau co-occurs with essentially the same mitochondrial haplotype as *w*Ma that infects *D. simulans* on Madagascar and elsewhere in Africa and the South Pacific (Rousset and Solignac 1995; Merçot and Poinsot 1998; Ballard 2000a; James and Ballard 2000; James et al. 2002; Ballard 2004), suggesting that *w*Mau and *w*Ma may be the same strain infecting different host species following introgressive *Wolbachia* transfer (see below). *w*Mau and *w*Ma phenotypes are also more similar to one another than to *w*No, with only certain crosses between *w*Ma-infected *D. simulans* males and uninfected *D. simulans* females inducing CI (James and Ballard 2000). Polymorphism in the strength of CI induced by *w*Ma could result from host modification of *Wolbachia*-induced CI (Reynolds and Hoffmann 2002; Cooper et al. 2017), or from *Wolbachia* titer variation that influences the strength of CI and/or the strength of CI rescue by infected females. Alternatively, the single-base-pair deletion in the *cinB*^*w*No^ homolog or other mutations that influence CI strength, could be polymorphic in *w*Ma. *w*Ma infection frequencies in *D. simulans* are intermediate on Madagascar (infection frequency = 0.25, binomial confidence intervals: 0.14, 0.40; James and Ballard 2000), consistent with no CI, suggesting replication of rarely observed *w*Ma CI is needed. Including *D. simulans* from the island of Réunion in this infection-frequency further supports the conjecture that *w*Ma causes little or no CI (infection frequency = 0.31, binomial confidence intervals: 0.20, 0.45; James and Ballard 2000). Unfortunately, no Madagascar *D. simulans* stocks available at the National *Drosophila* Species Stock Center were *w*Ma infected, precluding detailed analysis of this strain.

Our genomic data indicate that *w*Mau may maintain an ability to rescue CI, as the *cinA*^*w*No^ homolog is intact in *w*Mau genomes with only one nonsynonymous substitution relative to *cinA*^*w*No^; *cidA* in *w*Mel was recently shown to underlie transgenic-CI rescue (Shropshire et al. 2018). *w*Ma seems to sometimes rescue CI, but conflicting patterns have been found, and additional experiments are needed to resolve this (Rousset and Solignac 1995; Bourtzis et al. 1998; Merçot and Poinsot 1998; James and Ballard 2000; Merçot and Poinsot 2003; Zabalou et al. 2008). Future work that tests for CI rescue by *w*Mau and *w*Ma-infected females crossed with males infected with *w*No or other CI-causing strains, combined with genomic analysis of CI loci in *w*Ma, will be useful.

### *w*Mau does not influence *D. mauritiana* fecundity

While selection does not directly act on the level of CI (Prout 1994; Turelli 1994; Haygood and Turelli 2009), it does act to increase the product *Wolbachia*-infected host fitness and the efficiency of maternal transmission (Turelli 1994). Understanding the *Wolbachia* effects that lead to spread from low frequencies and the persistence of non-CI causing *Wolbachia* at intermediate frequencies is crucial to explaining *Wolbachia* prevalence among insects and other arthropods. The four-fold fecundity effect of *w*Mau reported by Fast et al. (2011) in *D. mauritiana* is inconsistent with our experiments and with the intermediate infection frequencies observed in nature. We find no *w*Mau effects on host fecundity, regardless of host background, with our estimates of *F* having BCa intervals that include 1. Small increases in *F* could allow the deterministic spread of *w*Mau from low frequencies, although detecting very small increases in *F* is difficult (Supplemental Figure 1B). Our results are consistent with an earlier analysis that assessed egg lay of a single genotype and found no effect of *w*Mau on host fecundity (Giordano et al. 1995). When combined with the low observed infection frequencies, our fecundity data are also consistent with our mathematical analyses indicating that *Wolbachia* can increase host fitness by at most about 50% for reasonable estimates of *μ*. Because fecundity is one of many fitness components, analysis of other candidate phenotypes for aiding the spread of low-frequency *Wolbachia* is needed.

### Introgressive *Wolbachia* transfer likely predominates in the *D. simulans* clade

Hybridization and introgression in the *D. simulans* clade may have led to introgressive transfer of *Wolbachia* among host species (Rousset and Solignac 1995). This has been observed in other *Drosophila* (Turelli et al. 2018; Cooper et al. 2019) and *Nasonia* wasps (Raychoudhury et al. 2009). The number of *Wolbachia* strains in the *D. simulans* clade, and the diversity of mitochondria they co-occur with, is complex. Figure 5A shows host relationships and Figure 5B shows mitochondrial relationships, with co-occurring *Wolbachia* variants in parentheses. While *D. mauritiana* is singly infected by *w*Mau, *D. simulans* is infected by several strains, including CI-causing *w*Ha and *w*No that often co-occur as double infections within individuals (O’Neill and Karr 1990; Merçot et al. 1995; Rousset and Solignac 1995). *w*Ha and *w*No are similar to *w*Sh and *w*Sn, respectively, that infect *D. sechellia* (Giordano et al. 1995; Rousset and Solignac 1995). *w*Ha and *w*Sh also occur as single infections in *D. simulans* and in *D. sechellia*, respectively (Rousset and Solignac 1995). In contrast, *w*No almost always co-occurs with *w*Ha in doubly infected *D. simulans* individuals (James et al. 2002), and *w*Sn seems to occur only with *w*Sh (Rousset and Solignac 1995). *D. simulans* has three distinct mitochondrial haplotypes (*si*I, *si*II, *si*III) associated with *w*Au/*w*Ri (*si*II), *w*Ha/*w*No (*si*I), and *w*Ma (*si*III). The *si*I haplotype is closely related to the *se* haplotype found with *w*Sh and *w*Sn in *D. sechellia* (Ballard 2000b). *w*Ma co-occurs with the *si*III haplotype, which differs over its 13 protein-coding genes by only a single-base pair from the *ma*I mitochondrial haplotype carried by *w*Mau-infected *D. mauritiana*. A second haplotype (*ma*II) is carried by only uninfected *D. mauritiana* (Ballard 2000a; James and Ballard 2000).

The lack of whole *w*Ma genome data precludes us from confidently resolving the mode of *w*Mau acquisition in *D. mauritiana*. However, mitochondrial relationships support the proposal of Ballard (2000b) that *D. mauritiana* acquired *w*Mau and the *ma*I mitochondrial haplotype via introgression from *w*Ma-infected *D. simulans* carrying *si*III. *D. mauritiana* mitochondria are paraphyletic relative to *D. sechellia* and *D. simulans* mitochondria (Solignac and Monnerot 1986; Satta and Takahata 1990; Ballard 2000a, 2000b), with *ma*I sister to *si*III and *ma*II outgroup to all other *D. simulans*-clade haplotypes (see Figure 5). Of the nine genomes we assessed, all but one (uninfected-*R44*) carry the *ma*I haplotype, and genotypes carrying *ma*I are both *w*Mau-infected (*N* = 5) and uninfected (*N* = 3). While *w*Ma-infected *D. simulans* carry *si*III, *w*No-infected *D. simulans* carry *si*I. We estimate that *w*Mau and *w*No diverged about 55,000 years ago, with only 0.068% sequence divergence over 682,494 bp. Nevertheless, it seems implausible that *w*No (versus *w*Ma) was transferred directly to *D. mauritiana* as this requires horizontal or paternal transmission of *w*No into a *D. mauritiana* background already carrying the *ma*I mitochondrial haplotype. Although our nuclear result suggests a confident phylogenetic resolution of the *D. simulans* clade (Figure 5A), this is an artifact of the bifurcation structure imposed by the phylogenetic analysis. Population genetic analyses show a complex history of introgression and probable shared ancestral polymorphisms (Kliman et al. 2000) among these three species. Consistent with this, of the 20 nuclear loci we examined, 6 (*aconitase, aldolase, bicoid, ebony, enolase, ninaE*) supported *D. mauritiana* as the outgroup within the *D. simulans* clade, 7 (*glyp, pepck, pgm, pic, ptc, transaldolase, wingless*) supported *D. sechellia* as the outgroup, and 7 (*esc, g6pdh, glys, pgi, tpi, white, yellow*) supported *D. simulans*. With successive invasions of the islands and purely allopatric speciation, we expect the outgroup to be the island endemic that diverged first. Figure 5B indicates that the *ma*II haplotype diverged from the other mtDNA haplotypes roughly when the clade diverged, with the other haplotypes subject to a complex history of introgression and *Wolbachia*-associated sweeps, as described by Ballard (2000b).

Ballard (2000b) estimated that *si*III-*ma*I diverged about 4,500 years ago, which presumably approximates the date of the acquisition of *w*Mau (and *si*III, which became *ma*I) by *D. mauritiana*. This is surely many thousands of generations given previous estimates that consider the temperature dependence of *Drosophila* development (Cooper et al. 2014; Cooper et al. 2018). As shown by our mathematical analyses (Eq. 3), the apparent persistence of the *ma*II mtDNA among *Wolbachia*-uninfected *D. mauritiana––*without its occurrence among infected individuals*––*is unexpected. More extensive sampling of natural *D. mauritiana* populations is needed to see if this unexpected pattern persists. The persistence of this haplotype is inconsistent with simple null models, possibly indicating interesting fitness effects.

While paternal transmission has been observed in *D. simulans* (Hoffmann and Turelli 1988; Turelli and Hoffmann 1995), it seems to be very rare (Richardson et al. 2012; Turelli et al. 2018). *w*No almost always occurs in *D. simulans* individuals also infected with *w*Ha, complicating this scenario further. It is possible that horizontal or paternal transmission of *w*Ma or *w*No between *D. simulans* backgrounds carrying different mitochondrial haplotypes underlies the similarities of these strains within *D. simulans*, despite their co-occurrence with distinct mitochondria. Given the diversity of *Wolbachia* that infect *D. simulans*-clade hosts, and known patterns of hybridization and introgression among hosts (Garrigan et al. 2012; Brand et al. 2013; Garrigan et al. 2014; Matute and Ayroles 2014; Schrider et al. 2018), determining relationships among these *Wolbachia* and how *D. mauritiana* acquired *w*Mau will require detailed phylogenomic analysis of nuclear, mitochondrial, and *Wolbachia* genomes in the *D. simulans* clade.

## AUTHOR CONTRIBUTIONS

MM performed the molecular and phenotypic work, participated in the design of the study, and contributed to the writing; WC performed the phylogenetic and genomic analyses and contributed to the writing; SR contributed to the molecular and phenotypic analyses and to the writing; JB performed the library preparation and contributed to the writing; MT contributed to the analyses, data interpretation, and writing; BSC designed and coordinated the study, contributed to the analyses and data interpretation, and drafted the manuscript. All authors gave final approval for publication.

## ACKNOWLEDGMENTS

We thank Margarita Womack for sampling the *D. mauritiana* used in this study and Daniel Matute for sharing them. We thank Michael Gerth for sharing *Nomada* genomic data. Isaac Humble, Maria Kirby, and Tim Wheeler assisted with data collection. Michael Gerth, Michael Hague, and Amelia Lindsey provided comments that improved earlier drafts of this manuscript. Computational resources were provided by the University of Montana Genomics Core. Research reported in this publication was supported by the National Institute Of General Medical Sciences of the National Institutes of Health (NIH) under Award Number R35GM124701 to B.S.C. The content is solely the responsibility of the authors and does not necessarily represent the official views of the NIH. The authors declare no conflicts of interest.

## Appendix 1. Mathematical analyses of mtDNA and *Wolbachia* dynamics

Our analysis follows the framework developed in Turelli et al. (1992), but is simplified by the lack of CI. We suppose that introgression introduces a cytoplasm carrying *Wolbachia* and a novel mtDNA haplotype, denoted B. Before *Wolbachia* introduction, we assume the population is monomorphic for mtDNA haplotype A. With imperfect maternal *Wolbachia* transmission, uninfected individuals will be produced with mtDNA haplotype B. Without horizontal or paternal transmission (which are very rare, Turelli et al. 2018), all *Wolbachia*-infected individuals will carry mtDNA haplotype B. Once *Wolbachia* is introduced, uninfected individuals can have mtDNA haplotype A or B. We assume that these three cytoplasmic types (“cytotypes”) differ only in fecundity, and denote their respective fecundities *F*_I_, *F*_A_ and *F*_B_. Denote the frequencies of the three cytotypes among adults in generation t by *p*_I,t_, *p*_A,t_ and *p*_B,t_, with *p*_I,t_ + *p*_A,t_ + *p*_B,t_ = 1. Without CI, the frequency dynamics are

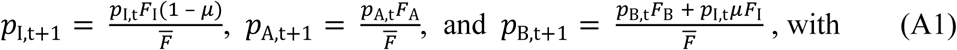

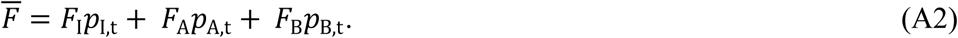

If the uninfected population is initially monomorphic for mtDNA haplotype A, the *Wolbachia* infection frequency will increase when rare if and only if

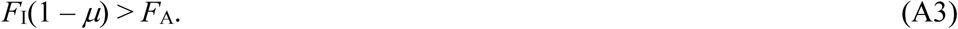

Turelli et al. (1992) showed that if a CI-causing *Wolbachia* is introduced with a cytoplasm that contains a novel mtDNA haplotype B, which has no effect on fitness, *Wolbachia*-uninfected individuals will eventually all carry haplotype B. This follows because eventually all uninfected individuals have *Wolbachia*-infected maternal ancestors. This remains true for non-CI-causing *Wolbachia* that satisfy (A3). However, we conjectured that if the introduced B mtDNA is deleterious in the new host nuclear background, i.e., *F*_A_ > *F*_B_, a stable polymorphism might be maintained for the alternative mtDNA haplotypes. The motivation was that imperfect maternal transmission seemed analogous to migration introducing a deleterious allele into an “island” of uninfected individuals. To refute this conjecture, consider the equilibria of (A1) with

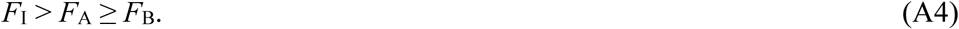

If all three cytotypes are to be stably maintained, we expect each to increase in frequency when rare. In particular, we expect the fitness-enhancing mtDNA haplotype A to increase when the population contains only infected individuals and uninfected individuals carrying the deleterious *Wolbachia*-associated mtDNA haplotype B. From (A1), *p*_A,t_ increases when rare if and only if

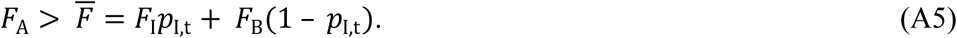

In the absence of haplotype A, we expect *p*_I_ to be at equilibrium between selection and imperfect maternal transmission, i.e.,

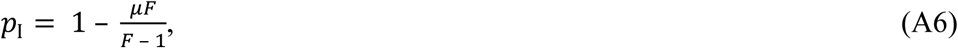

with *F* = *F*_I_ /*F*_B_ (Hoffmann and Turelli 1997). Substituting (A6) into (A5) and simplifying, the condition for *p*_A,t_ to increase when rare is

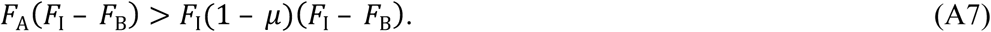

By assumption (A4), *F*_I_ > *F*_B_; hence (A7) contradicts condition (A3), required for initial *Wolbachia* invasion. Thus, simple selection on *Wolbachia*-uninfected mtDNA haplotypes cannot stably maintain an mtDNA polymorphism. The “ancestral” mtDNA haplotype A is expected to be replaced by the less-fit *Wolbachia*-associated haplotype B.

To understand the time scale over which this replacement occurs, let *r*_t_ denote the frequency of haplotype A among *Wolbachia*-uninfected individuals, i.e., *r*_t_ = *p*_A,t_ /(*p*_A,t_ + *p*_B,t_). From (A1),

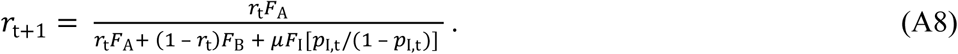

If we assume that the mtDNA haplotypes do not affect fitness, i.e., *F*_A_ = *F*_B_, and that the *Wolbachia* infection frequency has reached the equilibrium described by (A6), (A8) reduces to

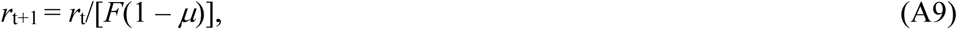

with *F* = *F*_I_ /*F*_B_.

## SUPPLEMENTAL INFORMATION

## Supplemental Tables

**Supplemental Table 1.** *w*Mau assembly statistics.

| **Supplemental Table 1.** *w*Mau assembly statistics. |  |  |  |  |
| --- | --- | --- | --- | --- |
| **Genotype** | **Scaffold count** | **N50 (bp)** | **Longest scaffold (bp)** | **Total length (bp)** |
| *R9* | 36 | 60,027 | 169,305 | 1,266,004 |
| *R29* | 36 | 61,106 | 169,295 | 1,277,467 |
| *R31* | 39 | 63,676 | 169,381 | 1,272,847 |
| *R41* | 42 | 61,106 | 170,537 | 1,303,156 |
| *R60* | 38 | 63,156 | 221,751 | 1,282,564 |

**Supplemental Table 2.**
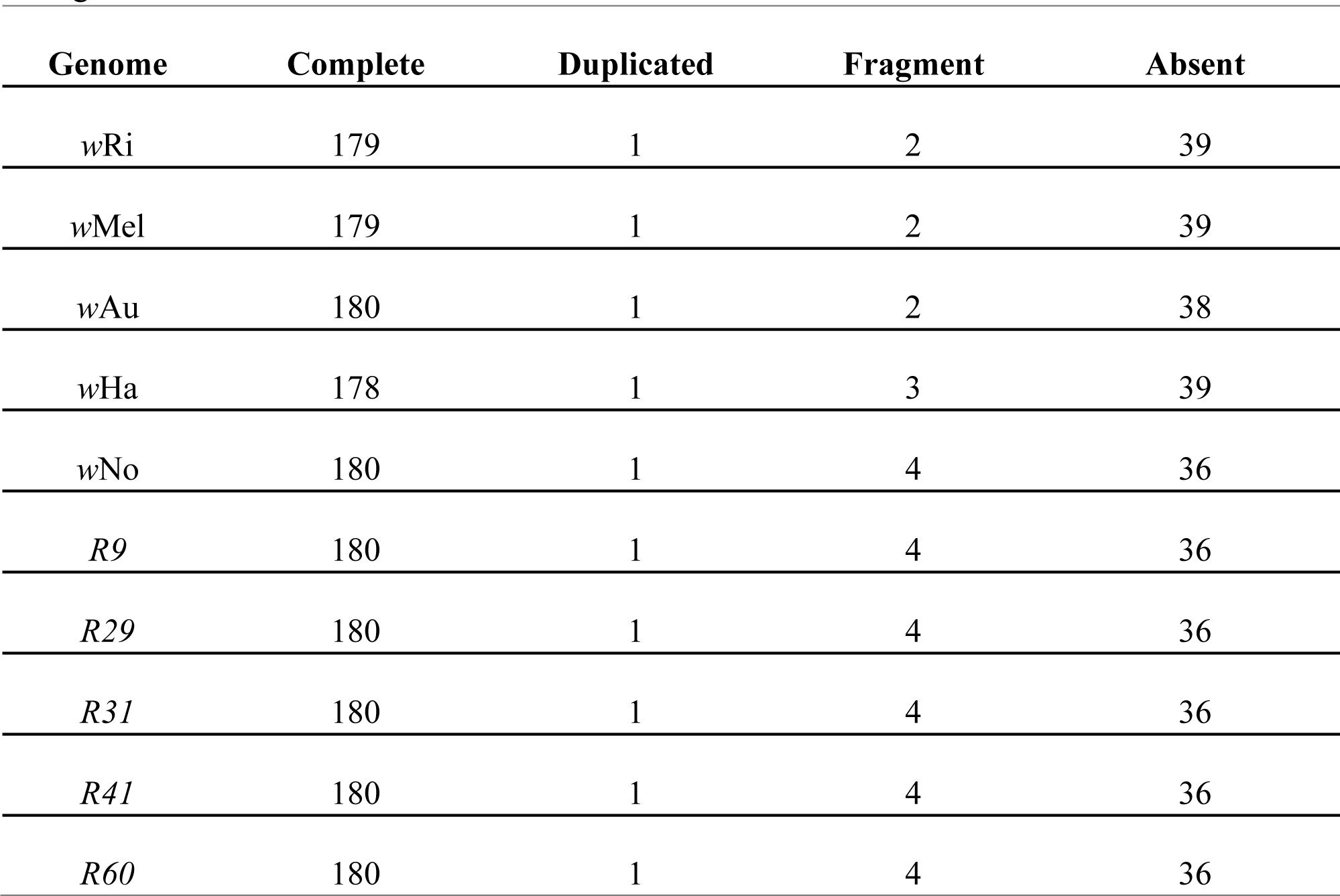
Near-universal, single-copy proteobacteria genes (out of 221) found using BUSCO v. 3.0.0.

**Supplemental Table 3.** Copy number variants in *w*Mau relative to sister *w*No. Genomic positions are based on the *w*No reference. There were no CNVs among *w*Mau variants.

| Start | End | Change | Wilcoxon Rank Sum Test | Kolmogorov-Smirnov Test |
| --- | --- | --- | --- | --- |
| 115,000 | 142,000 | 1 → 0 | < 0.0001 | < 0.0001 |
| 590,000 | 593,000 | 1 → 0 | 0.0027 | 0.0050 |
| 982,000 | 991,000 | 1 → 0 | < 0.0001 | < 0.0001 |
| 1,195,000 | 1,199,000 | 1 → 0 | 0.0005 | 0.0007 |

**Supplemental Table 4:** Genes present in regions deleted in *w*Mau relative to *w*No. Genes predicted to be pseudogenized in *w*No are shaded grey.

| Accession number | Name |
| --- | --- |
| **Deletion 1 (115,000-142,000):** |  |
| wNO_RS00550 | Hypothetical protein |
| wNO_RS06015 | Ankyrin repeat domain protein |
| wNO_RS00560 | Pseudo IS256 family transposase, frameshifted |
| wNO_RS00565 | Recombinase family protein |
| wNO_RS00570 | DUF2924 domain-containing protein |
| wNO_RS00575 | Ankyrin repeat domain-containing protein |
| wNO_RS00580 | Ankyrin repeat domain-containing protein |
| wNO_RS00585 | Phage tail protein |
| wNO_RS00590 | Baseplate assembly protein GpJ |
| wNO_RS00595 | Pseudo baseplate assembly protein W, frameshifted |
| wNO_RS00600 | Hypothetical protein |
| wNO_RS00605 | Phage baseplate assembly protein V |
| wNO_RS00610 | Hypothetical protein |
| wNO_RS00615 | Putative minor tail protein Z |
| wNO_RS00620 | Hypothetical protein |
| wNO_RS00625 | Minor capsid protein E |
| wNO_RS00630 | Head decoration protein |
| wNO_RS00635 | S49 family peptidase |
| wNO_RS00640 | Pseudo phage portal protein, frameshifted |
| wNO_RS00645 | Phage head stabilizing protein GpW |
| wNO_RS00650 | Phage terminase large subunit family protein |
| wNO_RS00655 | Ankyrin repeat domain-containing protein |
| wNO_RS00660 | Hypothetical protein |
| wNO_RS00665 | Hypothetical protein |
| wNO_RS00670 | Sigma-70 family RNA polymerase sigma factor |
| wNO_RS00675 | ATP-binding protein |
| wNO_RS00680 | IS110 family transposase |
| **Deletion 2 (590,000-593,000):** |  |
| wNO_RS02645 | XRE family transcriptional regulator |
| wNO_RS02650 | Hypothetical protein |
| wNO_RS02655 | Hypothetical protein |
| wNO_RS02660 | XRE family transcriptional regulator |
| **Deletion 3 (982,000-991,000):** |  |
| wNO_RS04690 | Group II intron reverse transcriptase/maturase |
| wNO_RS06355 | Hypothetical protein |
| wNO_RS04695 | Pseudo hypothetical protein, partial |
| wNO_RS04700 | Pseudo cell filamentation protein Fic, partial |
| wNO_RS04705 | Hypothetical protein |
| wNO_RS04710 | DNA methylase |
| wNO_RS04715 | Hypothetical protein |
| wNO_RS04720 | Ankyrin repeat domain-containing protein |
| wNO_RS04725 | Phage terminase large subunit family protein |
| wNO_RS04730 | Phage head stabilizing protein GpW |
| wNO_RS04735 | DUF1016 domain-containing protein |
| **Deletion 4 (1,195,000-1,199,000):** |  |
| wNO_RS06095 | Ankyrin repeat domain protein |
| wNO_RS05590 | Rpn family recombination-promoting nuclease/putative transposase |

**Supplemental Table 5.**
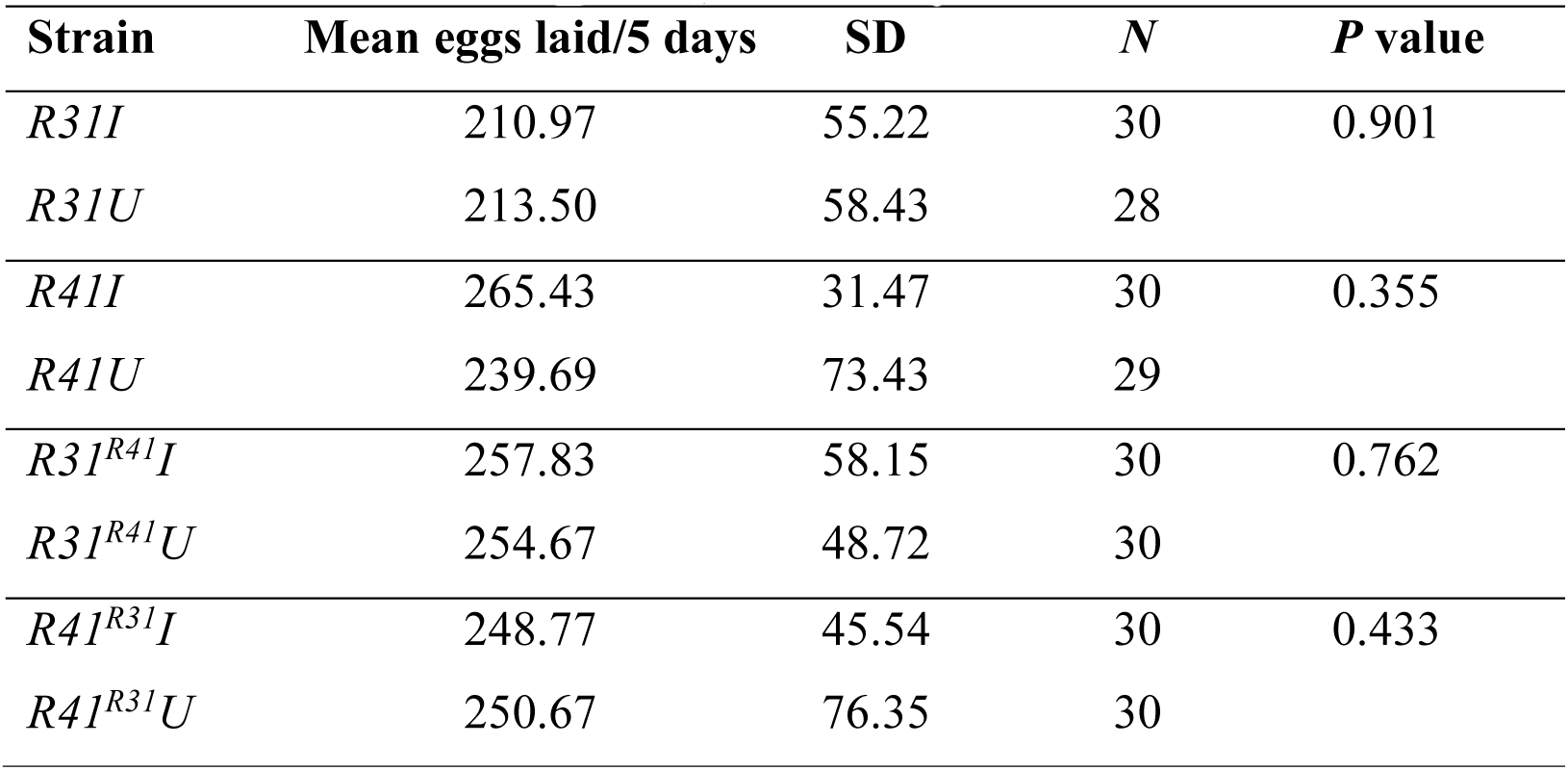
*w*Mau does not significantly affect *D. mauritiana* fecundity in comparisons of paired infected (I) and uninfected (U) strains sharing host nuclear backgrounds. *N* is the number of females that produced the means and SDs. *P* values are for two-tailed Wilcoxon tests. Cytoplasm sources are denoted with superscripts for introgressed strains.

## Supplemental Figures

**Supplemental Figure 1.**
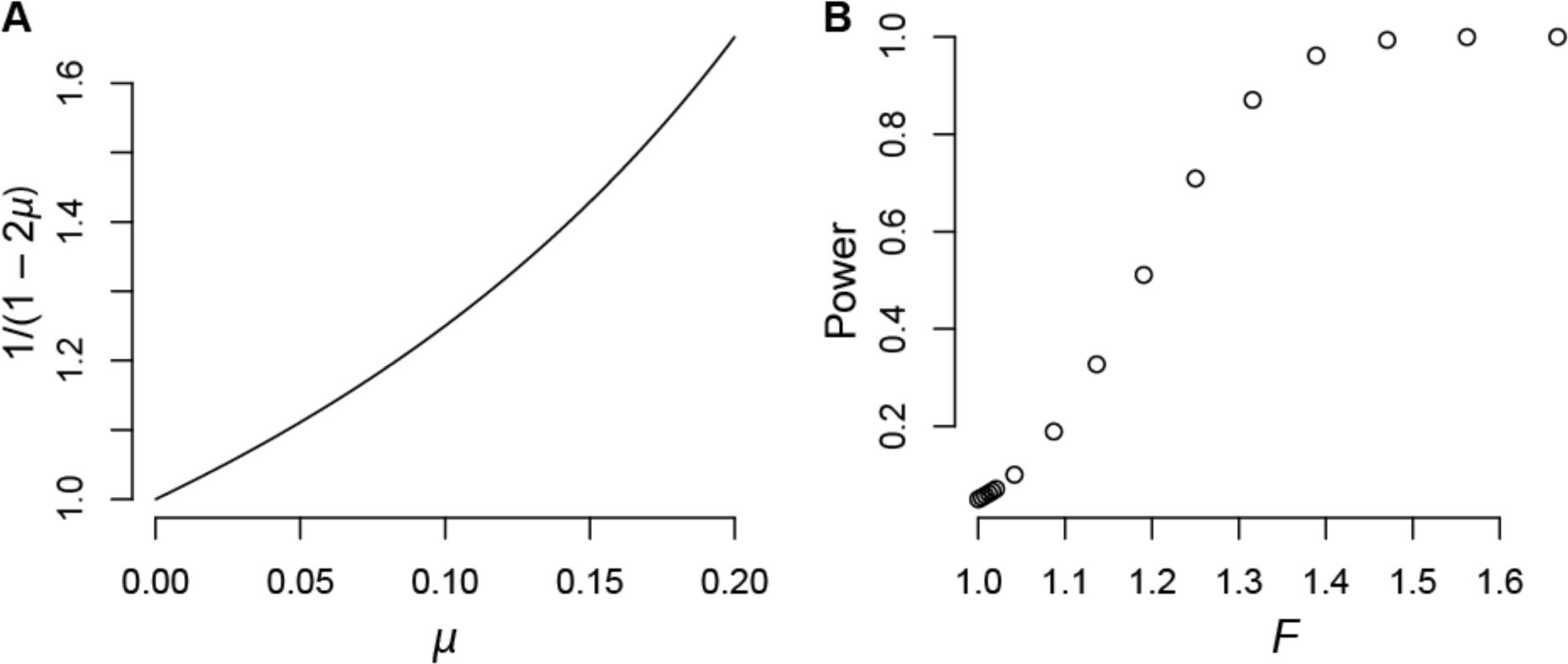
A) Our proposed upper bound, 1/(1 – 2*μ*), on plausible relative fitness, *F*, as a function of *μ* ranging from perfect transmission (*μ* = 0) to a level of imperfect transmission (*μ* = 0.2) that has not been observed in nature. B) Our powers to detect values of *F* with α = 0.05.

